# Engineering synthetic CopT/A-based genetic circuits for miRNA imaging and functional gene regulation

**DOI:** 10.1101/2023.01.25.525516

**Authors:** Chuanxian Zhang, Xiaorui Shi, Chu Tang, Yarong Du, Wenjie Shu, Fu Wang

## Abstract

Synthetic genetic circuits that can operate at the transcriptional and translation levels have been widely applied in the control of cellular behaviors and functions. However, the regulation of genetic circuits is often accompanied by the introduction of exogenous substances or the endogenous generation of inhibitory products, which would bring uncontrollable hazards for biological safety and reduce the efficiency of the system. Here, we described a miRNA-responsive CopT-CopA (miCop) genetic circuit system to realize real-time monitoring the intracellular expression of miRNA-124a during neurogenesis or miRNA-122 under the stimulation of extracellular drugs in living cells and animals. Furthermore, to prove the modularity of the system, we engineered this mipCop to tune the expression of DTA (diphtheria toxin A) gene, and showed the powerful capacity of killing cancer cells by inducing apoptosis and cell cycle arrest based on miRNA response. This study provides an effective means to couple miRNA sensing with miRNA-responsive gene modulation, which may open up new diagnostic or therapeutic applications.

## Introduction

MicroRNAs (miRNAs) are a class of small noncoding RNAs of approximately 22 nucleotides that regulate gene expression at the transcriptional or post-transcriptional level^1^. MiRNAs interact sequence-specifically with the 3’UTR of target gene to induce mRNA degradation or inhibit protein translation, thus acting as gene regulators^2^. It is estimated that miRNAs can mediate more than 30% of the human genome. Since miRNAs play critical roles in various biological processes such as cell proliferation, apoptosis, inheritance and metabolism, they are evolutionarily conserved and differentially expressed in various organisms^3^. Moreover, miRNAs are frequently dysregulated in a range of malignant cancers. Some overexpressed miRNAs act as oncogenes, while some downregulated ones act as tumor suppressor genes, indicating potential biomarkers for cancer diagnosis^4 5^. Thus, there is urgent need to develop tools for detecting and visualizing the dynamic expression of miRNAs in mammalian cells and animals.

As a multidisciplinary biotechnology, synthetic biology employs various components to assemble and construct genetic circuits, which can exert various biological functions in a variety of environments, especially in the precise diagnosis and treatment of diseases. Synthetic bioluminescence reporters, which include a variety of enzymes (such as luciferase) and substrates (D-uciferin, etc)^6-9^, have attracted extensive interests for miRNA imaging in diverse biological processes. Bioluminescence itself does not depend on the absorption of light by any organism, and the biochemical reaction of luciferase with luciferin substrates releases luminescence with high sensitivity^10^. In addition, synthetic and endogenous miRNAs have been widely applied in combination with other regulatory elements, such as the CRISPR-Cas12a-designed synthetic gene circuit loop system, to expand the regulatory topology^11^. By designing “miRNA switch gene circuits”, cell populations can be clearly separated based on the expression of reporter genes, the risk of genome damage can be effectively reduced^12^, and the precise identification of trace miRNAs in human serum samples can also be achieved^13^.There are also studies demonstrating that repeated administration of nanoparticles containing miRNA switches significantly reduces neointima formation after wire injury and allows vessel reendothelialization. Furthermore, this cell-selective nano-therapy is a valuable tool with the potential to advance the fight against neointimal hyperplasia and atherosclerosis^14^. The miRNA-based classifier synthetic gene circuit design builds a framework for preclinical classifier research, combining animal models, delivery platforms, in vivo expression levels of inputs, and efficacy of outputs, provides an effective evaluation for the preclinical application of synthetic gene circuits^15^. However, most of the current synthetic gene circuits targeting miRNA are often controlled by the introduction of exogenous substances, such as ribozyme-based riboswitches^16^, macromolecular proteins^17^, etc., or endogenous production of inhibitory products^18^, and the structure is also more complex, which poses a great threat to biological safety and reduces the efficiency of synthesizing gene circuits.

To address these limitations, we attempted to develop a miRNA-responsive CopT-CopA (miCop) system to achieve efficient gene circuit engineering with high targeting and biosafety. CopA is the master control element of E. coli R1 plasmid replication^19^. It is an antisense RNA of approximatively 90 nt in length with stem-loop structure. It is transcribed from a complementary strand in the leader region of repA mRNA. Another complementary sequence is CopT, which can form a complex with CopA in the form of kissing^20^. The translation of the genes between the CopT-CopA will be hindered due to the steric hindrance effect^21^. These unique functions make it an excellent gene regulatory tool. Therefore, in this study, we engineered CopT-CopA into a genetic circuit system, realizing *in vitro* blood detection and *in vivo* imaging of miRNAs in living cells and animals. Furthermore, we evaluated the specific anti-tumor effect of this miCop reporter in tumor cells through activating a cytotoxic fragment A of diphtheria toxin (DTA). Our results provide a promising genetic strategy for miRNA detection and gene regulation, which may open up new diagnostic or therapeutic applications.

## Results and Discussions

### Rational design of miCop

In order to develop an efficient miRNA-responsive system with high targeting and biosafety, we constructed the miCop biosensor using the CopT-CopA gene assembly, which does not require the addition of exogenous substances, nor does it endogenously generate by-products. To facilitate the miRNA-guided cleavage, three copies of fully complementary miRNA target sequences (3xTargets) were inserted into the 3’untranslational region (3’UTR) of gene of interest (GOI). An internal ploy(A) was immediately placed after the GOI to protect the transcript from degradation. When miRNAs are absent in this system, CopA can form a complex with CopT in the form of kissing, resulting in a steric hindrance effect to hinder the translation of the GOI that are between them. In the case of miRNA present in cells, the target miRNA would bind to the 3xTargets sequences, thereby cleaving the downstream CopA sequence and releasing the translation suppression, enabling the expression of intermediate functional gene elements (Fig. 1).

**Figure 1.**
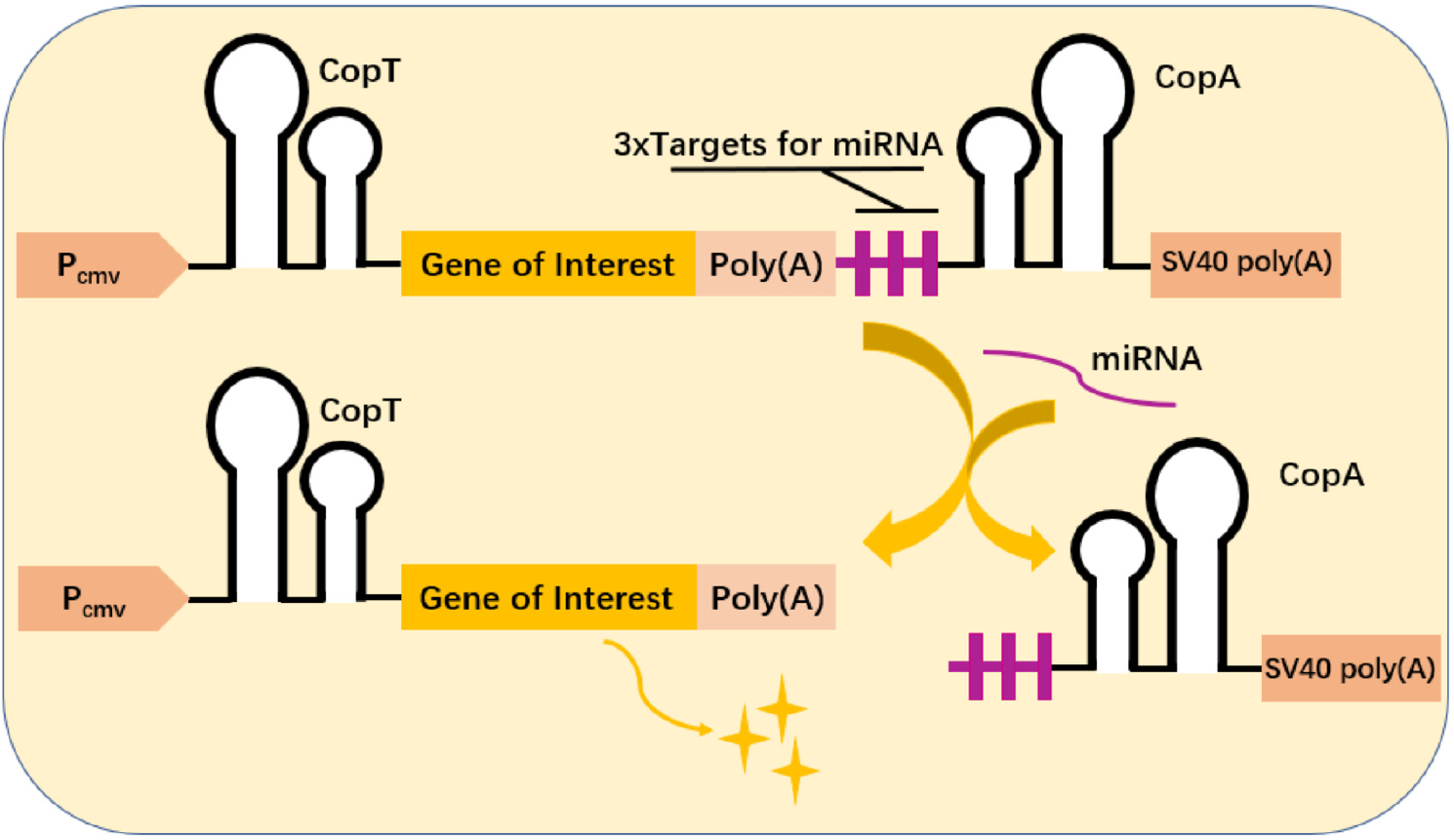
Schematic representation of the miCop system. When the target miRNAs are present, they will be completely complementary to the target sequences of the miRNAs, thereby excising the downstream CopA sequences, releasing the inhibition of the CopT-CopA complex, and allowing the expression of the functional gene element between CopT and CopA. P_cmv_, CMV promoter.

### Characterization of miCop

To characterize the miCop system, we first selected firefly luciferase (Fluc) as a gene of interest to verify the function of CopT-CopA gene assembly. The Fluc gene was inserted between CopT and CopA to generate Cop-Fluc reporter. We also constructed a control Fluc reporter without CopT-CopA pairs (Fig. 2A). The three plasmids Cop-Fluc, Fluc, and Vector (empty) were transiently transfected into Hela (Fig. 2B), 293 (Fig. 2C) or P19 cells (Fig. 2D), respectively. Then the luciferase activity from each group was examined. The results showed that the luciferase signal in Cop-Fluc-transfected cells was significantly reduced compared with the Fluc group, indicating CopT-CopA effectively inhibited the protein expression of Fluc gene. However, the Fluc mRNA levels had no obvious change between Fluc and Cop-Fluc-transfected cells (Fig. 2E). These results confirmed that the association of CopT and CopA would form a second structure and prevent the translation but not transcription of the intervening Fluc gene.

**Figure 2.**
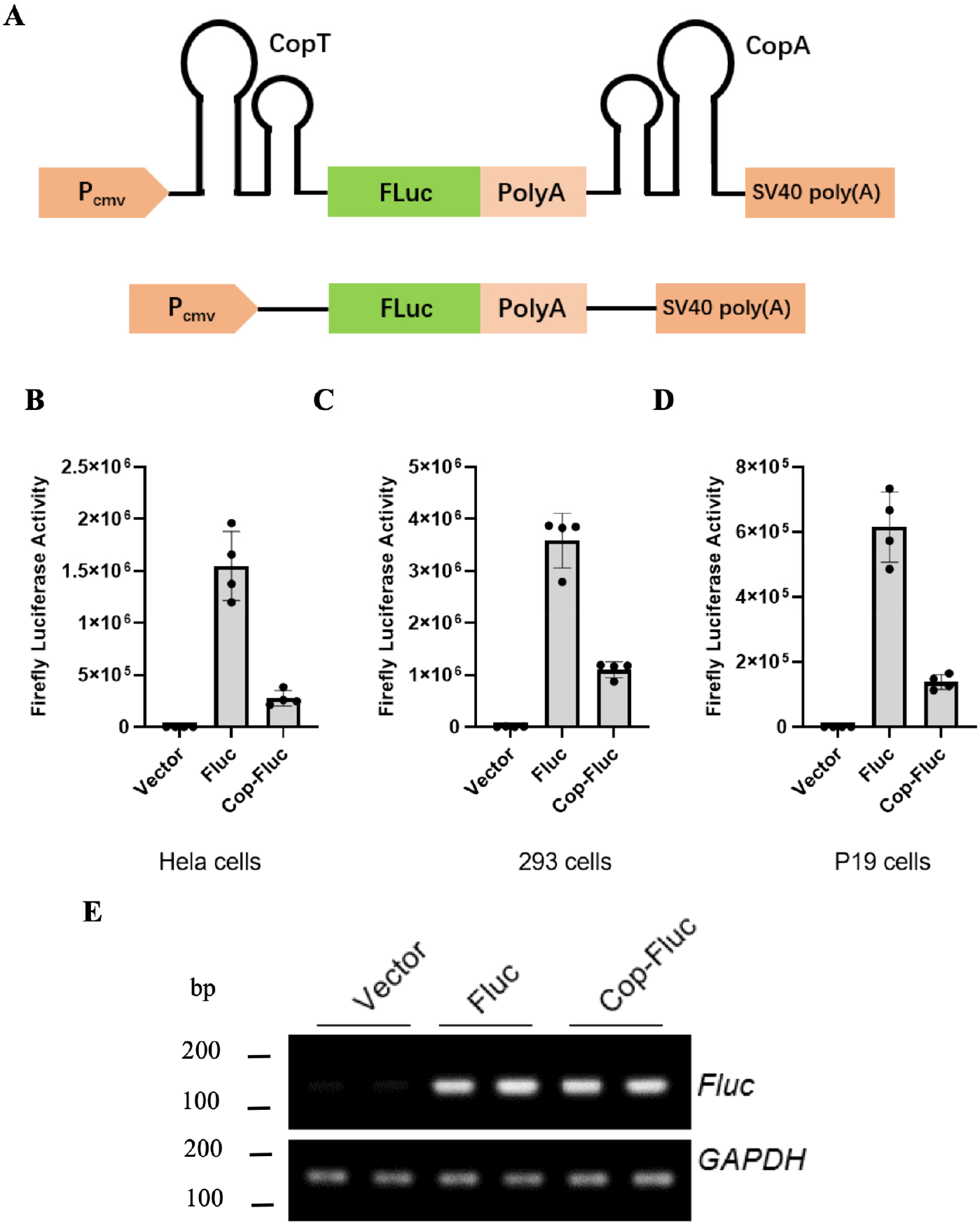
Functional validation of the CopT-CopA gene assembly. (A) Fluc was inserted as a functional gene between CopT-CopA to construct a Cop-Fluc reporter. A control Fluc reporter was also constructed without CopT-CopA pairs. (B, C, D) The inhibitory effect of the CopT-CopA gene assembly was verified by detecting the expression level of luciferase gene. The Vector (empty), Fluc and Cop-Fluc were transfected into (B) Hela cells, (C) 293 cells, and (D) P19 cells, respectively. The luciferase activity of Fluc gene was measured in each group. (E) RT-PCR assay was performed to detect the Fluc mRNA levels in the Vector (empty), Fluc or Cop-Fluc transfected Hela cells.

After validation of the repression effect of CopT-CopA, we then test whether the reporter transcript could be cleaved by miRNA. Three copies of target sequences for miRNA-124a, a neuro-specific miRNA, were cloned upstream of CopA to obtain the miCop-Fluc biosensor (Fig. 3A). The Fluc activity in 293 cells exhibited a dose-dependent increase upon treatment with exogenous miRNA-124a mimics, indicating that the Fluc signal was liberated by miRNA-124a-dependent cleavage (Fig. 3B). We then explored the signal response to the different concentrations of miCop-Fluc reporter with a fixed amount of miRNA-124a (40 nM). The results demonstrated that the Fluc signal raised according to the increased dose of transfected miCop-Fluc plasmids (Fig. 3C). Therefore, we determined the optimal concentration of miRNA-124a (80 nM) and miCop-Fluc reporter (20 ng) for subsequent experiments (Fig. 3D). In addition, the luciferase activity could be activated only in the presence of exogenous miRNA-124a but not miRNA-1 or miRNA-133, indicating the cleavage specificity of miCop-Fluc reporter by miRNAs (Fig. 3E).

**Figure 3.**
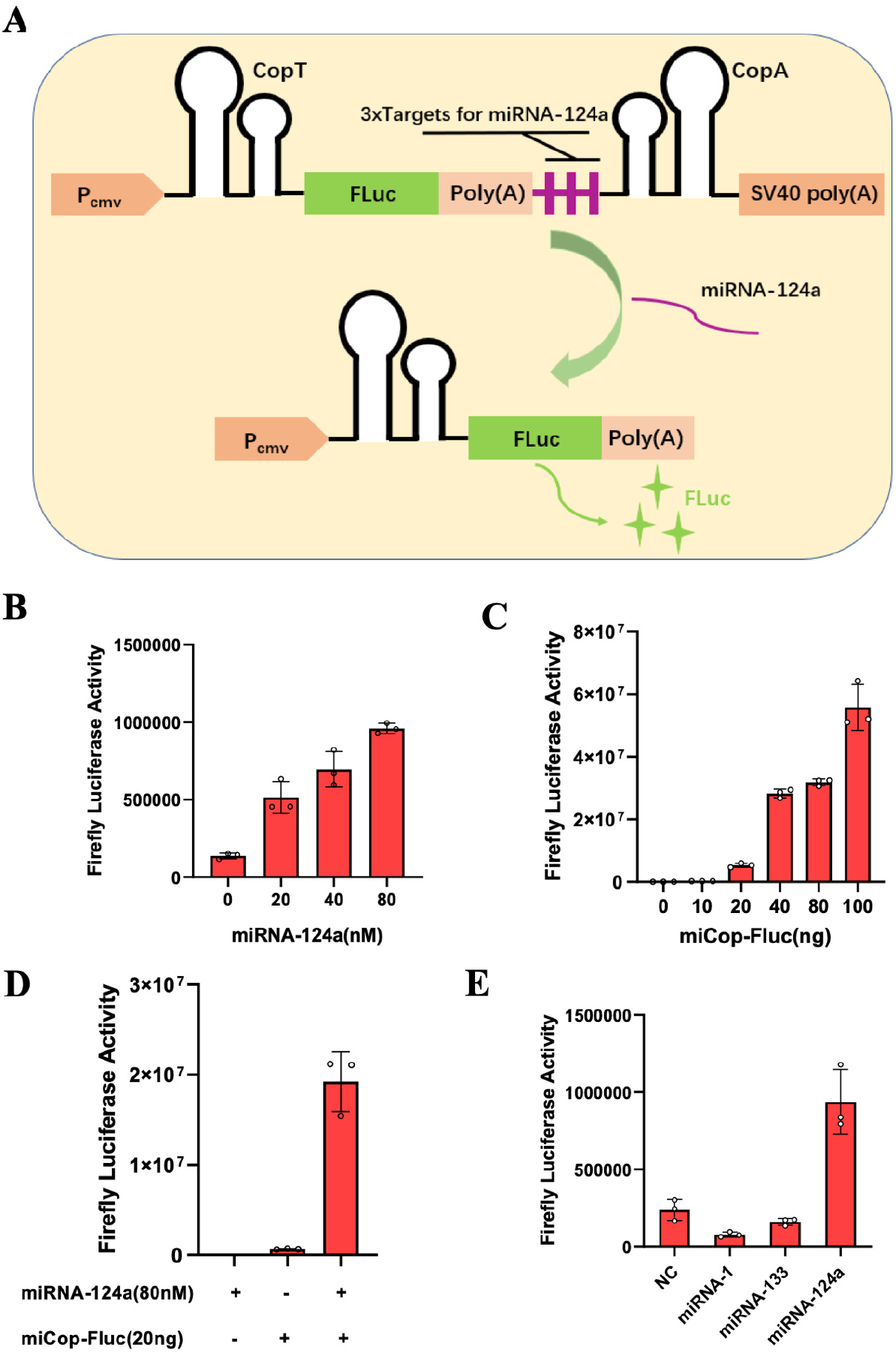
Development and characterization of miCop-Fluc. **(A)** Schematic representation of the miCop-Fluc. When miRNA-124a is present, it binds to the target sequence and cleaves the downstream CopA sequence, leading to the Fluc expression. (B) Different concentrations (0, 20, 40, 80 nM) of miRNA-124a mimics and miCop-Fluc were co-transfected into 293 cells, and the luciferase activity was measured 24 h later. (C) Different amounts (0, 10, 20, 40, 80 ng) of miCop-Fluc and miRNA-124a mimics (40 nM) were co-transfected into 293 cells, and the luciferase activity was measured. (D) 20 ng miCop-Fluc and 80 nM miRNA-124a were selected as the optimal concentrations for verification. (E) Different miRNA mimics (NC, miRNA-1, 133, 124a) and miCop-Fluc were co-transfected into 293 cells, and the luciferase activity was measured.

### Real-time imaging of miRNA-124a in neurogenesis with miCop-Fluc

To explore the feasibility of this miCop-Fluc reporter for monitoring the endogenous miRNA expression, a neurogenesis cellular model was established using P19 cells. P19 cells are a pluripotent stem cell line isolated from mouse teratoma, and can differentiate into neurons or glial cells under the induction of retinoic acid^22,23^. The expression level of miRNA-124a in neural precursor cells is very low, but the expression gradually increases during neural development and reaches a peak after neural cell maturation^24^. Therefore, we used retinoic acid to induce the differentiation of P19 cells for 0, 2, 4, and 6 days, respectively. The bright images showed that with the increase of induction time, P19 cells gradually transformed into neuronal morphology, and the number of protrusions gradually increased and lengthened (Supplementary Fig. S1). The confocal images demonstrated the upregulation of neuronal marker MAP2 over time in P19 cells, confirming the successful differentiation of P19 cells towards neuronal cells (Fig. 4A).

**Figure 4.**
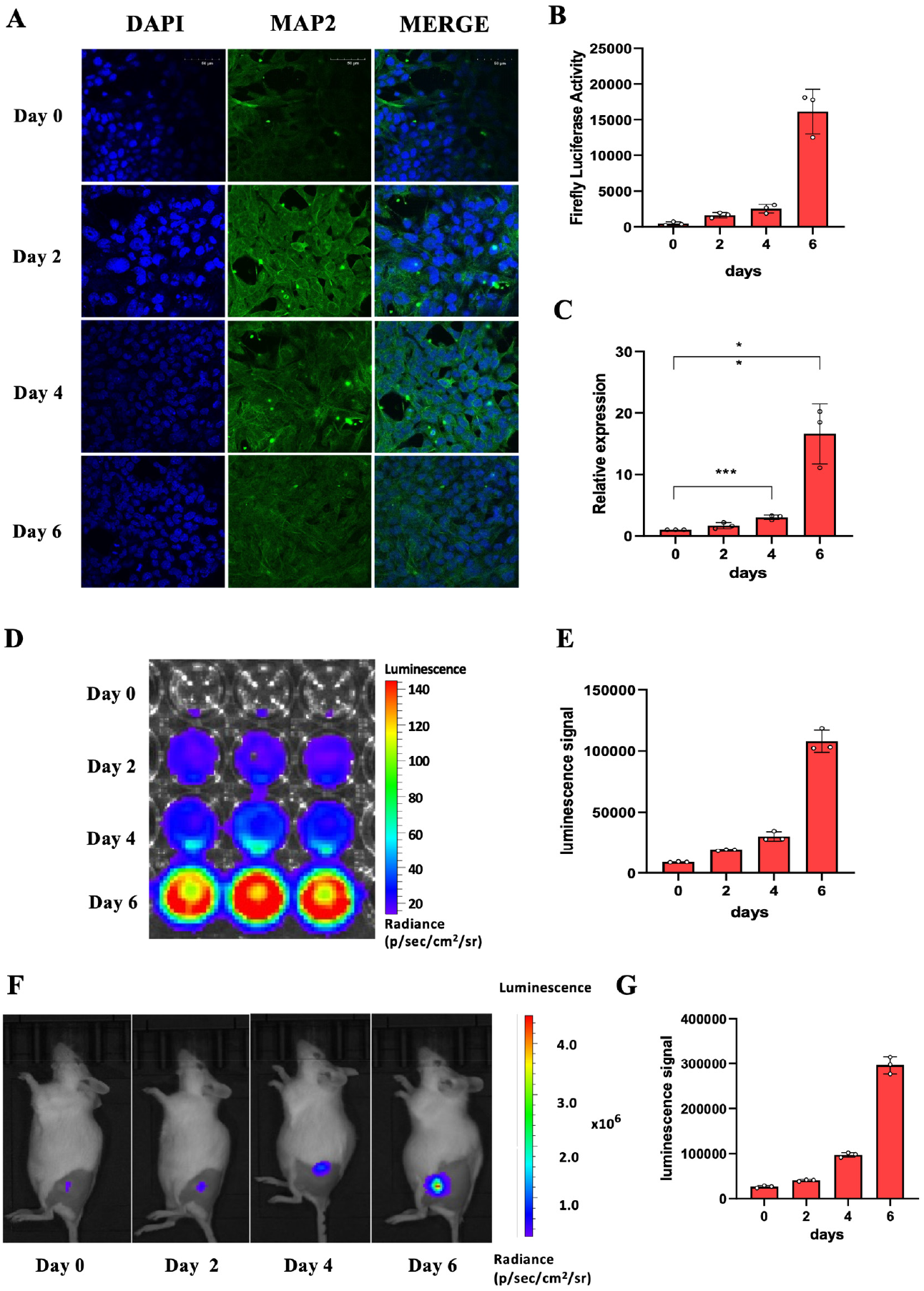
Real-time visualiztion of miRNA-124a in neurogenesis with miCop-Fluc. (A) Immunofluorescence analysis of MAP2 expression after retinoic acid-induced differentiation of P19 cells in day 0, 2, 4, 6. (B) The miCop-Fluc was transfected into differentiated P19 cells for different days, and their luciferase activities were measured. (C) qPCR was performed to detect the expression level of miRNA-124a in differentiated P19 cells at different days. (D) The miCop-Fluc was transfected into differentiated P19 cells for different days, and the bioluminescence signal was captured. (E) The quantification of the bioluminescence signals in (D). (F) The miCop-Fluc-transfected P19 cells were injected subcutaneously into the legs of mice. Then the mice were treated with retinoic acid to induce cell differentiation for different days. Finally, the bioluminescence intensity of the mice was captured. Representative images were shown. (G) The quantification of the bioluminescence signals in (F).

After establishing the neurogenesis cellular model, miCop-Fluc reporter was transfected into the differentiated P19 cells to monitor the endogenous miRNA-124a dynamic expression during neurogenesis. After the induction by retinoic acid for different days (0, 2, 4, 6 day), the luciferase signal gradually increased over time, representing the activated miR-124a expression (Fig. 4B). The qPCR data confirmed that the expression level of endogenous miRNA-124a increased during P19 neurogenesis (Fig. 4C). The *in vitro* bioluminescence signal of miCop-Fluc reporter was also found to respond to endogenous miRNA-124a during the process of retinoic acid treatment (Fig. 4D, E). These results suggested that our miCop-Fluc reporter could monitor the dynamic expression of endogenous miRNA-124a during neurogenesis *in vitro*. In order to explore the performance of miCop-Fluc for *in vivo* imaging of miRNAs, the P19 cells were transfected with miCop-Fluc plasmids and then implanted subcutaneously into living mice. Then the mice were treated with retinoic acid to induce cell differentiation. As shown in Figure 4F and 4G, a gradually increase of bioluminescence signal was observed over the induction process. Collectively, these results suggest that the miCop-Fluc biosensor enables noninvasively imaging of miRNA-124a in a real-time manner.

### Cellular monitoring of miR-122 with miCop-Gluc

To further explore the application of miCop reporter in monitoring miRNA activity, we engineered *Gaussia princeps* luciferase (Gluc), a secreted luciferase from the marine copepod *Gaussia princeps*, into the miCop system to examine the liver-specific miRNA-122 expression. Abnormal expression of miRNA-122 is associated with liver diseases^25,26^. MiRNA-122 levels were reduced in rodent and human-derived hepatocellular carcinoma compared with healthy liver^27^. The three copies of target sequences for miRNA-122 were inserted upstream of CopA to obtain miCop-Gluc biosensor (Fig. 5A). Due to the good secretory property, the miRNA activity can be evaluated by directly measuring Gluc signal in cell supernatants without splitting cells. To validate the miCop-Gluc biosensor, different concentrations of miRNA-122 mimics and miCop-Gluc constructs were transfected into 293T cells. As shown in Fig. 5B, the Gluc activity in cell supernatants increased according to the increased dose of exogenous miRNA-122, indicating the miRNA-122 activated the Gluc signal of miCop-Gluc (Fig. 5B). In order to explore the response of miCop-Gluc to endogenous miRNA-122, different amounts of miCop-Gluc plasmids were transfected into HepG2 cells, where miRNA-122 was intrinsically expressed. The results showed that the Gluc signal also increased in a dose dependent manner under the activation of endogenous miRNA-122 (Fig. 5C). Moreover, the Gluc activation is specific to miRNA-122 but not other miRNA such as miRNA-1, indicating the specificity of miCop-Gluc reporter in monitoring miRNAs (Fig. 5D).

**Figure 5.**
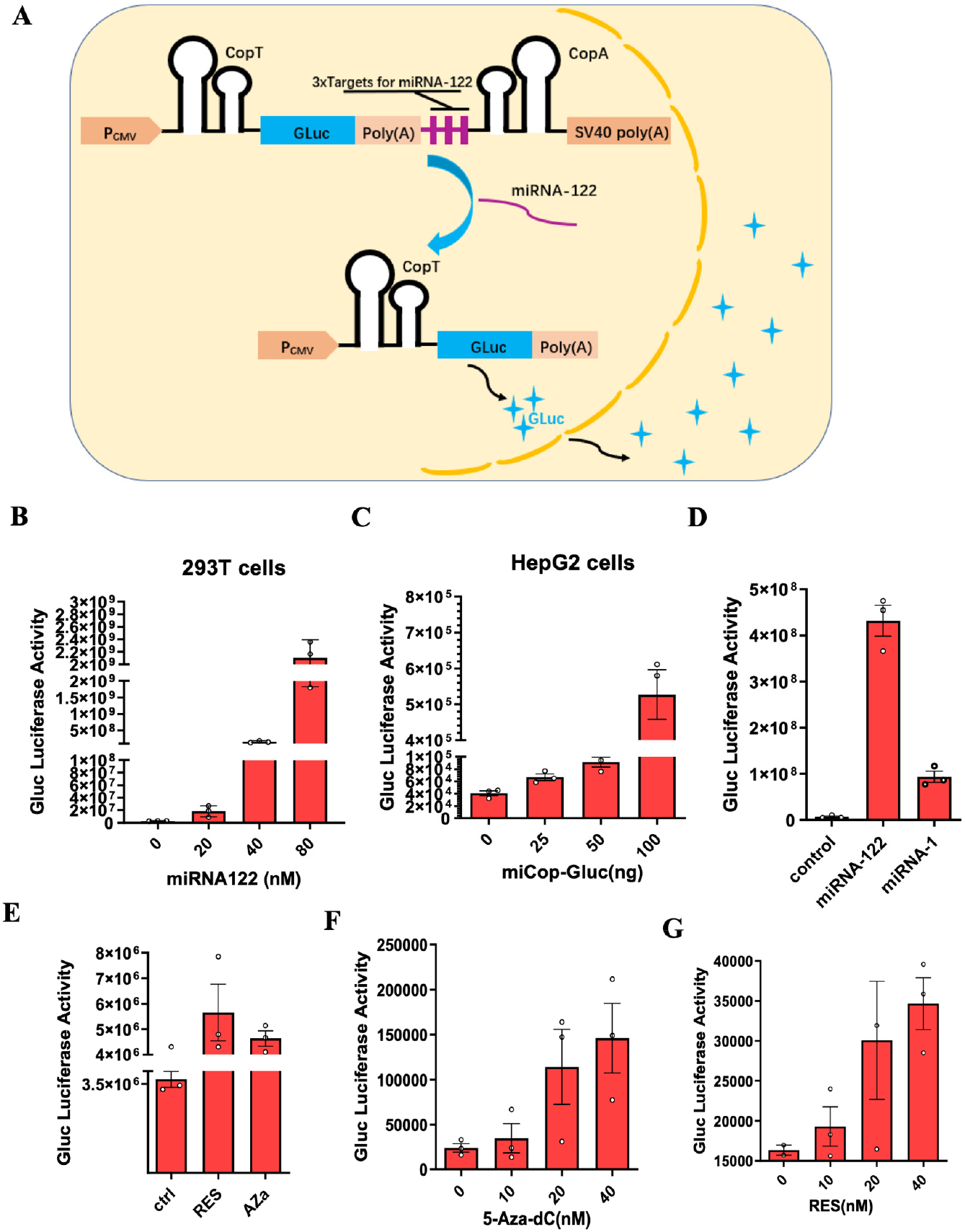
Development and characterization of a miCop-Gluc system. (A) Schematic representation of the miCop-Gluc. When miRNA-122 is present, it binds to the target sequence, thereby cleaves the downstream CopA sequence, leading to the Gluc expression. Gluc signal is secreted and detected in cell supernatants. (B) 293 cells were co-transfected with different concentrations of miRNA-122 mimics and miCop-Gluc plasmids, and Gluc activity was measured in cell supernatants. (C) The miCop-Gluc constructs at different concentrations were transfected into HepG2 cells to detect the response of the reporter to endogenous miRNA-122. The Gluc activity was measured in cell supernatants. (D) 293 cells were co-transfected with miCop-Gluc plasmids and miRNA-122 or miRNA-1 mimics, respectively. Cell supernatants were assayed for Gluc signal. (E) HepG2 cells were transfected with miCop-Gluc constructs and treated with resveratrol (RES) or 5-Aza-dC (AZa) for 24 h. Then the Gluc activity was measured in the cellular supernatant. (F, G) The miCop-Gluc-transfected HepG2 cells were treated with different concentrations of (F) 5-Aza-dC or (G) resveratrol or. 24 h later, the expression level of Gluc in the cellular supernatant was detected.

To further examine the dynamic expression changes of miRNA-122 induced by extracellular stimuli, two drugs, resveratrol (RES) and 5-aza-2-deoxycytidine (5-Aza-dc), were used to treat cells. Resveratrol, a polyphenolic compound, was found to chemosensitize doxorubicin-resistant cancer cells by modulating miRNA-122-5p^28^. 5-Aza-dc is a commonly used demethylating agent^29^, and the expression of miRNA-122 was upregulated by 5-Aza-dc in HepG2 and Hep3B cells ^30,31^. As shown in Figure 5E, the Gluc signal was significantly increased in HepG2 cells after treatment with RES or 5-Aza-dc, indicating both drugs activated the expression of endogenous miRNA-122, which is consistent with the previous studies. Additionally, the signal increase showed a good dose-dependent (Fig. 5F, 5G). These results suggest that the miCop-Gluc can realize the monitoring of miRNA-122 expression changes triggered by exogenous drugs.

### *In vivo* imaging and *ex vivo* blood monitoring of miR-122 with miCop-Gluc

In order to explore the applicability of miCop-Gluc reporter for *in vivo* imaging of miRNA-122, 293 cells were transfected with miCop-Gluc and miR-122 mimics or NC, then the cells were collected and subcutaneously injected into both thighs of mice to establish 293 cells-implanted mouse model. As shown in Figure 6A and 6B, the luminescence signal in miR-122 group was significantly higher than that in NC group, indicating the feasibility of miCop-Gluc reporter for *in vivo* imaging. We then employed the miCop-Gluc reporter to monitor miRNA-122 expression after treatment with 5-Aza-dc. miCop-Gluc-transfected HepG2 cells were implanted subcutaneously in mice and then the both thighs side of mice were treated with 5-Aza-dC or control (Figure 6C). The luminescence signal was obviously increased after 5-Aza-dC treatment in the left side of mice (Figure 6D). Moreover, the signal on the left thigh of mice increased with the increase of 5-Aza-dC concentration (10, 20, 40, 80 nM), confirming that 5-Aza-dC upregulated endogenous miR-122 and activated the Gluc signal (Figure 6E, F). Finally, based on the characteristics of Gluc secretion, we intravenously injected the miCop-Gluc reporter into mice, and collected blood from the tail vein at different time periods to detect the Gluc signal in the blood. As shown in Figure 6G, the Gluc signal in blood increased with time and reached the peak at 6 h time point, and then decreased afterwards. The results showed that the miCop-Gluc achieved a dynamic response to endogenous miRNA-122 in living body, providing a powerful sensor for miRNA detection both *ex vivo* and *in vivo*.

**Figure 6.**
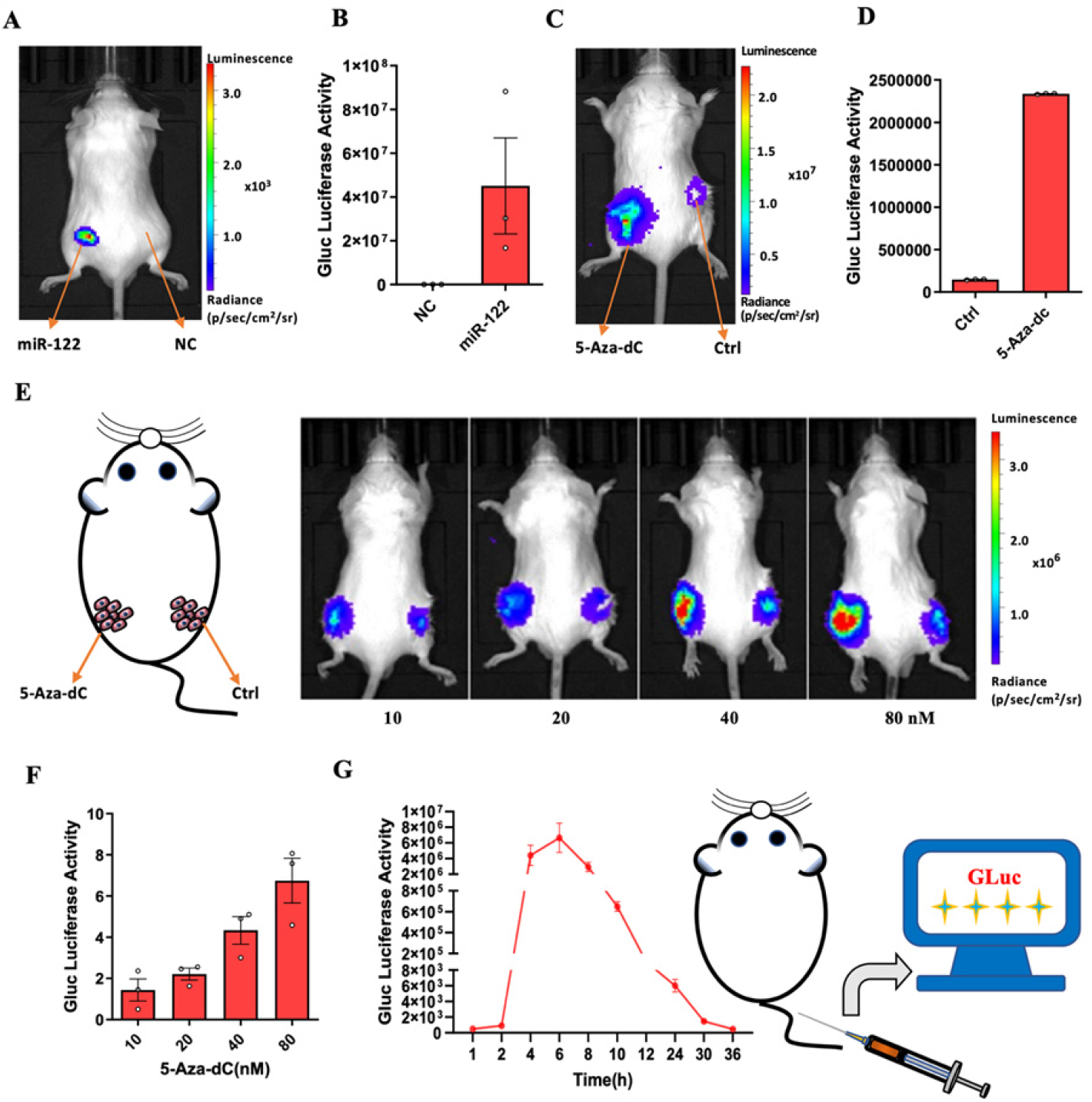
In vivo imaging and ex vivo blood monitoring of miR-122 activity. (A) The miCop-Gluc and miR-122 mimics or NC was co-transfected into 293 cells. 24 hours later, the cells were collected and injected into the both thighs of mice. Then the bioluminescence imaging was performed to capture the images. (B) Quantification of the in vivo imaging data. (C) HepG2 cells were transfected with miCop-Gluc plasmids and the cells were injected subcutaneously on the both sides of mice. The left and right thigh side of mice were treated with 5-Aza-dC or control. 24 h later, the bioluminescence imaging was performed. (D) Quantification of the in vivo imaging data. (E) miCop-Gluc-transfected HepG2 cells were implanted subcutaneously in both sides of mice and then the left thigh side of mice were treated with different concentrations (10, 20, 40, 80 nM) of 5-Aza-dC. PBS control was injected into the right side of the mouse. 24 h later, the bioluminescence imaging was performed. (F) Quantification of the in vivo imaging data. (G) The miCop-Gluc plasmids were intravenously injected into mice, and blood was collected from tail vein at different time points to detect the Gluc levels in blood.

### miCop-DTA induced cell apoptosis and cell cycle arrest via activating DTA gene

In order to further expand the application of the miCop system, we chosen DTA gene as the functional gene element of the miCop system to construct the miCop-DTA genetic circuit (Fig. 7A). Diphtheria toxin (DT) is a bacterial toxin secreted by pathogenic strains of *Corynebacterium diphtheriae*. Fragment A of DT (DTA) has a strong cytotoxic effect^32,33^, showing great potential in anticancer therapy. First, we transfected different concentrations of miRNA-124a mimics with a fixed amount of miCop-DTA vector into A549 cells, and performed the CCK-8 assay 24 hours later. The results showed that the cell viability decreased according to the increased dose of miRNA-124a (Fig. 7B), indicating the miCop-DTA caused cytotoxicity via miRNA-124a-dependent cleavage and activation of DTA expression. Furthermore, the miCop-DTA exhibited a dose-dependence on killing cells when increased amounts of miCop-DTA plasmids was transfected into A549, MCF-7 or 4T1 cells (Fig. 7C, D, E). In order to further verify the miCop-DTA system responds to endogenous miRNA-124a, P19 cells were differentiated with retinoic acid treatment for different days. The results showed that when the cell differentiation reached the fourth day, the killing effect of miCop-DTA on P19 cells had exceeded 80%, demonstrating the powerful anticancer ability of the miCop-DTA (Fig. 7F).

**Fig. 7.**
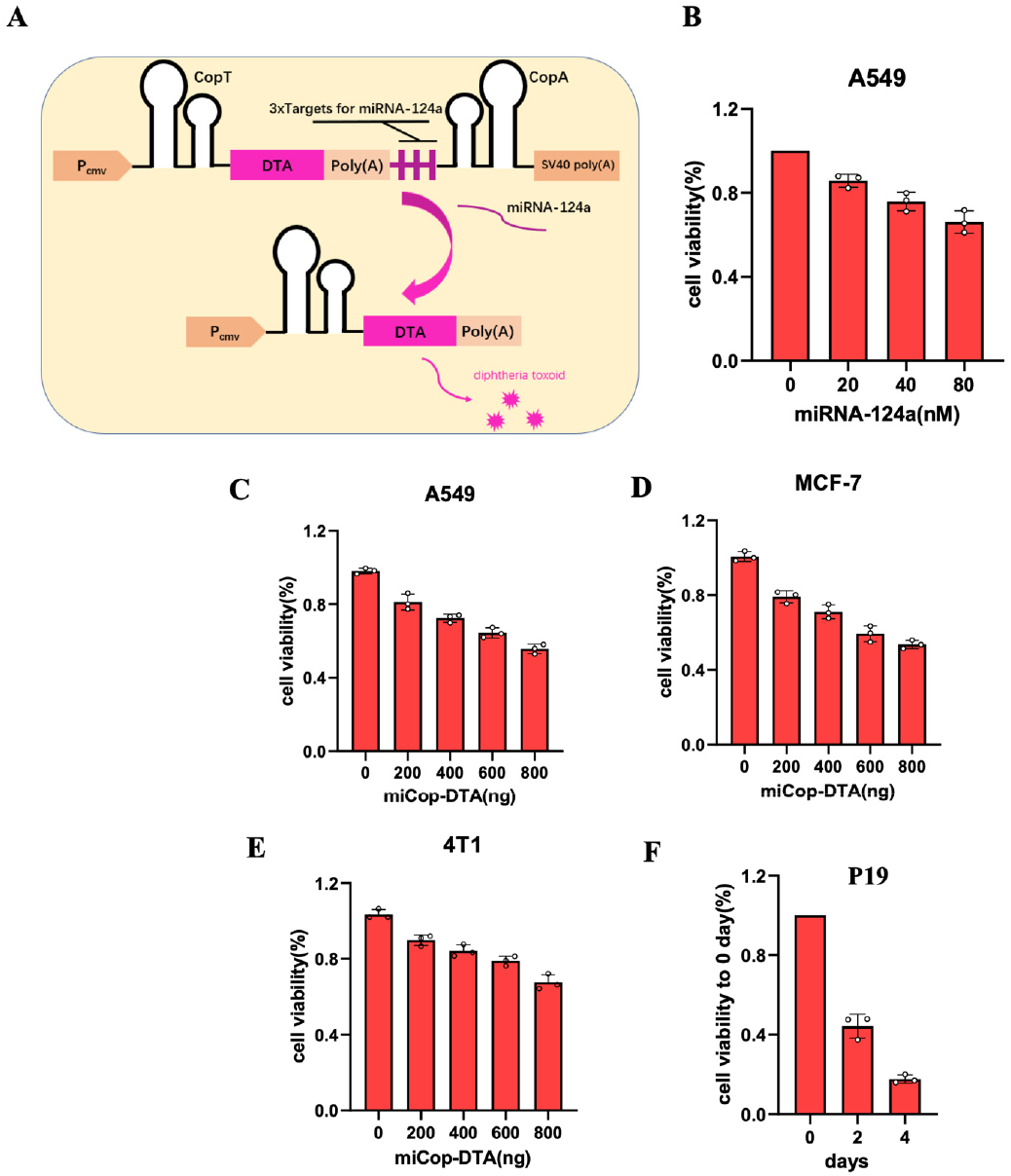
Design and characterization of miCop-DTA. (A) Schematic representation of the miCop-DTA. When miRNA-124a is present, it binds to the target sequence, thereby excising the downstream CopA sequence, activating the DTA gene, producing diphtheria toxin, and killing tumor cells. (B) Different concentrations of miRNA-124a mimics and with a fixed amount of miCop-Gluc were co-transfected into A549 cells, and cell viability was detected by CCK-8 assay 48 h later. (C-E) Different amounts of miCop-DTA plasmids was co-transfected with miRNA-124a mimics into (C) A549, (D) MCF-7 or (E) 4T1 cells. Then cell viability was detected by CCK-8 assay 48 h later. (F) P19 cells were differentiated with retinoic acid treatment for different days. Then miCop-Gluc plasmids were transfected into differentiated P19 cells. CCK-8 assay was performed to detect cell viability.

In order to explore whether miCop-DTA could affect the tumor cell migration, we performed wound healing assay in A549 cells. As shown in Fig. 8, the cells treatment with both miRNA-124a and miCop-DTA (miRNA-124a + miCop-DTA group) effectively inhibited the cell migration compared with the other treatment groups at different time points (Fig. 8). To further investigate the tumor-killing ability of miCop-DTA, flow cytometry was undertaken to detect the cell apoptosis. Similar to the wound healing assay, the PI/Annexin-V assay results showed that the miRNA-124a + miCop-DTA group induced a higher proportion of cell apoptosis than other groups (Fig. 9A). Next, we examined whether apoptosis was accompanied by cell cycle change. After treatment with both miRNA-124a and miCop-DTA, an obvious accumulation of cell cycle in S phase was observed (18.4% ± 1.7%) compared with the miRNA-124a or miCop-DTA alone group (Fig. 9B). Meanwhile, no significant changes in the G2/M phase were found among any of these five groups, whereas the G0/G1 phase population in miRNA-124a + miCop-DTA group was reduced to 60.1% ± 1.3%. These cell cycle findings are consistent with previous study^34^. Taken together, the results indicated that the intracellular activated DTA led to the arrested progression of cell cycle, which would further induce cell apoptosis.

**Figure 8.**
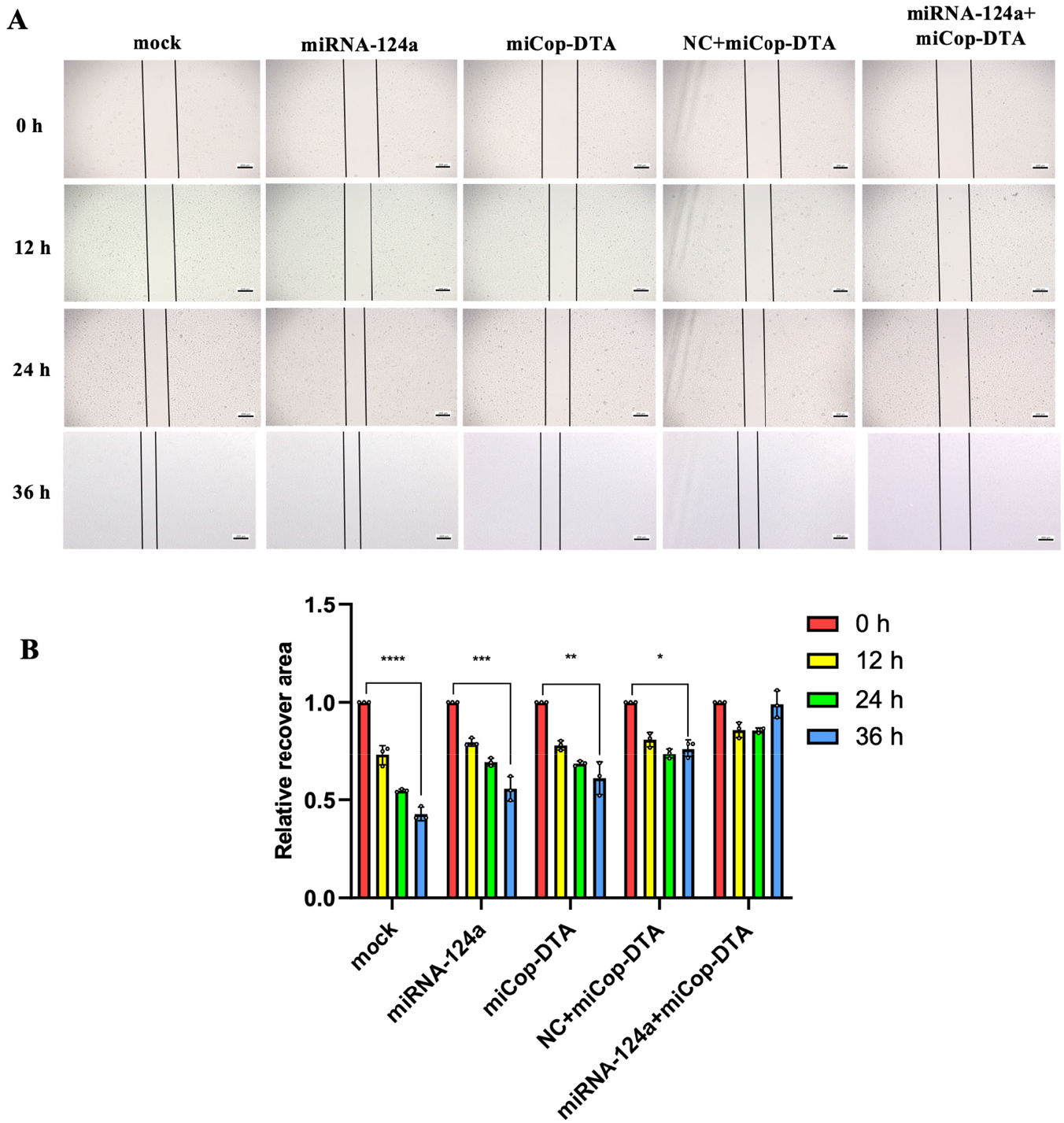
miCop-DTA inhibited tumor cell migration via activating DTA expression. (A) A549 cells were transfected with individual miRNA-124a, miCop-DTA, or both together. The wound healing was imaged at 0, 12, 24, and 36 hours using a microscope. The margins are outlined in black line (scale bar: 200 μm). (B) The recover areas were calculated in different treatment groups at different time point by using ImageJ software.

**Figure 9.**
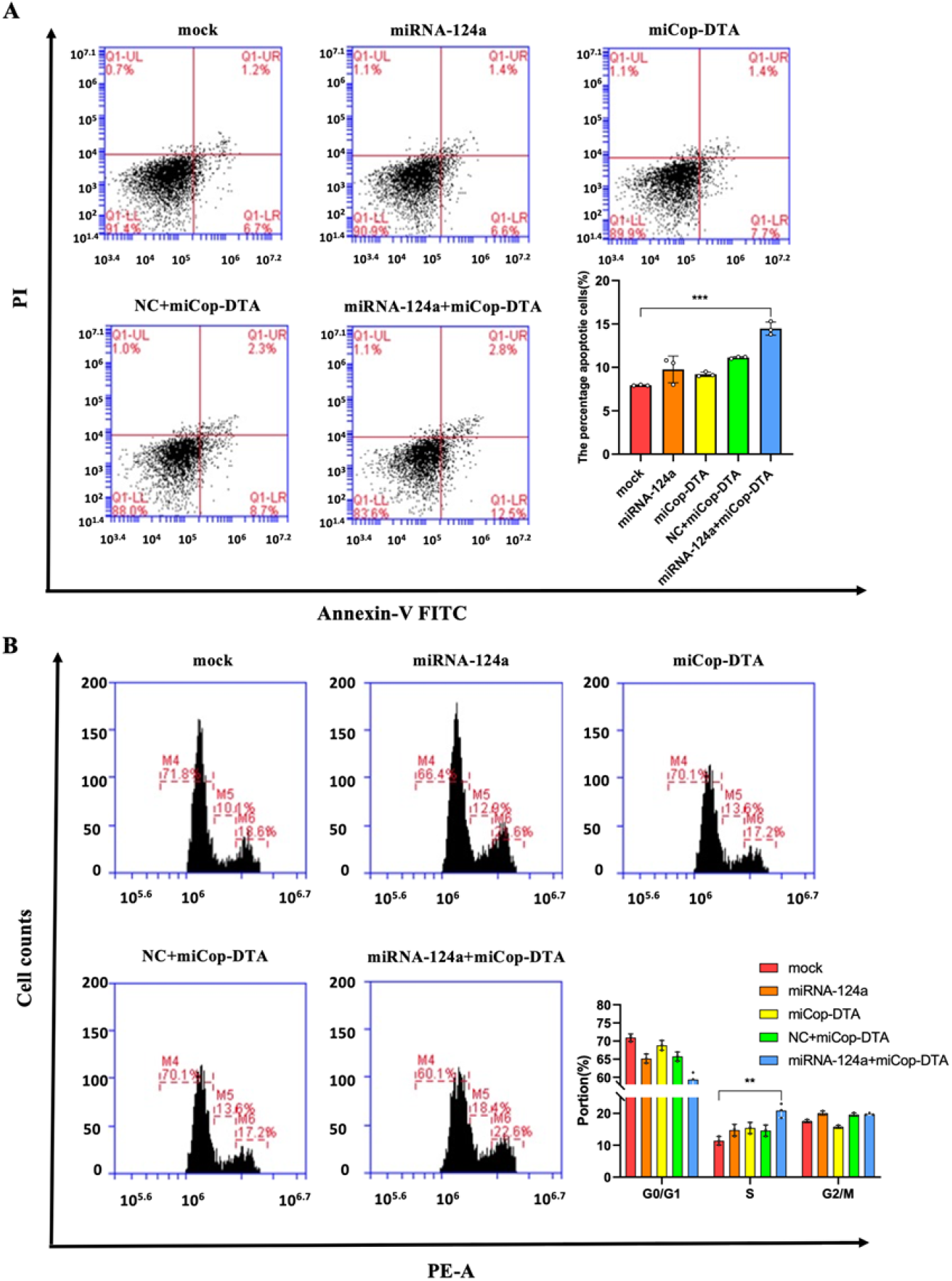
miCop-DTA induced cell apoptosis accompanied by cell cycle arrest. (A, B) A549 cells were transfected with individual miRNA-124a, miCop-DTA, or both together. The cells were collected and analyzed by (A) PI/AnnexinV FITC apoptosis assay or (B) cell cycle assay using flow cytometry. Three independent experiments were performed.

## Conclusion

In this study, we developed a CopT/A-based genetic engineering miCop system for miRNA imaging and gene activation. We demonstrated that miCop-Fluc and miCop-Gluc biosensors were capable of monitoring intracellular expression of miRNA-124a during neurogenesis or miRNA-122 under extracellular drug stimulation. In addition, by activating DTA gene expression, miCop-DTA showed the powerful capacity of killing cancer cells based on the miRNA levels. The ability to combine miRNA visualization with gene regulation enables our miR-Cop system a potential tool for RNA sense and RNA-responsive gene control.

## Materials and Methods

### Plasmid construction

To construct the miCop-Fluc plasmid, the oligonucleotides coding for CopT, Cop A, Fluc, or 3x miRNA Targets were synthesized (Shanghai Generay Biotechnology, Ltd), and cloned into the NheI and HindIII sites of the pcDNA3.1(+) vector. A 200 bp-long ploy(A) oligo was also synthesized and cloned between Fluc and 3x miRNA Targets sequences. To obtain the miCop-Gluc or miCop-DTA plasmid, Fluc sequence was replaced with the synthesized sequence for Gluc or DTA (Shanghai Generay Biotechnology, Ltd). Finally, all the recombinant plasmid was confirmed by DNA sequencing.

### Cell lines, reagents and transfection

The human cell lines 293, A549, MCF7, 4T1 and P19 were maintained in our lab. Cells were incubated in DMEM (Gibco, Carlsbad, CA, USA) supplemented with 10% fetal bovine serum (Gibco) and 100 U/mL penicillin/streptomycin (Gibco) in a humidified 5% CO2 incubator at 37 °C. The miRNA-124a or miRNA-122 mimics were synthesized by Shanghai GenePharma Ltd. For transfection, the cells were seeded in 24-well plates and cultured overnight. The next day, the plasmids and miRNA mimics were transfected into cells using Lipofectamine 2000 (Invitrogen, Carlsbad, CA, USA) according to the manufacturer’s instructions.

### Reverse Transcription PCR (RT-PCR) and quantitative PCR (qPCR)

To detect the mRNA expression level of Fluc, total RNA was isolated from cells using Trizol reagent (Invitrogen, Grand Island, NY). RNA samples (50 ng) were reverse transcribed using the QuantScript RT Kit (TianGen, Beijing) for cDNA synthesis. The PCR was performed in triplicate using PCR Mix (TianGen, Beijing) with Fluc primers at 94 °C for 5 min, 25 cycles of 94 °C 30 s, 56 °C 30 s and 72 °C 30 s. The PCR products were detected by agarose gel electrophoresis. GAPDH was used as loading control. To detect the miRNA-124a expression, the real-time PCR was performed using SYBR Green (miRNA qPCR Detection Kit, Biotake, China) at 94 °C for 5 min, 40 cycles of 94 °C for 30 s and 60 °C for 30 s with ABI 7700 PCR system (Applied Biosystems, USA). U6 snRNA was used as internal control. Each experiment was performed in triplicate. All the primer sequences are available upon request.

### P19 cell differentiation

Retinoic acid (RA) was prepared into a 5 μM stock solution using DMSO, and then diluted 1000 times into the prepared DMEM medium. The P19 cells were seeded in 24-well plates. After culturing for 24 hours, the cell medium was replaced with RA-supplemented medium every two days. After 6 days, P19 cells differentiated for 0, 2, 4, and 6 days were prepared.

### Confocal microscopy

P19 cells were treated with RA for differentiation for 0, 2, 4, and 6 days. Then the cells were washed with PBS for three times, each time for 5 minutes. The cells were then incubated with 4% cell fixation solution for 30 minutes at 4 ° C. The cells were then washed with PBS and incubated with DAPI solution and MAP2 FITC-labeled antibody. Live-cell imaging was taken using a laser confocal microscopy (Olympus, Japan) with Fluoview FV31S-SW Software (Olympus, Japan).

### Luciferase activity assay

Cells were plated in 24-well plates for 24 hours. The cells were transfected with different concentrations of plasmids and miRNA mimics for another 24 hours. After removal of the cell supernatant, the cells were washed with phosphate buffered saline (PBS, pH 7.4) and adherent cells were lysed with cell lysis buffer. For Fluc activity assay, 10 μL of cell lysate were mixed with 10 μL of D-luciferin substrate. The luciferase activity was measured using a Glomax-20/20 luminometer (Promega) according to the manufacturer’s instructions. For Gluc activity assay, 10 μL of cell supernatant was mixed with 10 μL of coelenterazine substrate. The Gluc luciferase activity was measured. Each experiment was performed in triplicate.

### CCK-8 assay

Cell Counting Kit (CCK)-8 was employed to determine the cell viability according to the manufacturer’s protocol (Beyotime Biotech Co. Ltd., Shanghai, China). Briefly, cells were seeded in a 96-well plate at the density of 5000 cells per well. After 24 h, the plasmids or miRNA mimics at different concentrations were transfected into cells for 24 h. Before determination, 10 μL CCK-8 solution was added to each well and incubated for another hour. Then the OD value of each well was measured at the filter of 450 nm through a microplate reader (Tecan Safire 2, Männedorf, Switzerland) and the cell viability was calculated. Each experiment was performed in triplicate.

### Wound healing Assay

A549 cells were seeded in 6-well plates and incubated until approximately 80% coverage was achieved. The plasmids or miRNA mimics were transfected into cells. Nest day, a sterile pipette tip was used to draw through the cell layer, then rinse twice with PBS to remove cellular debris. The cells were then incubated in a cell incubator and the wound closure was observed by microscopy at 0, 12, 24, and 36 hours. The relative recover area of wound healing was calculated with ImageJ software.

### PI/Annexin V-FITC apoptosis assay

A549 cells were seeded in 12-well plates. After 24 hours, the plasmids or miRNA mimics were transfected into cells and incubated for 24 hours. Then the cells were detached, washed and resuspended in PBS. The cells were double stained with propidium iodide (PI) and Annexin V-FITC kit (Beyotime Biotechnology Co., Ltd., Shanghai, China). The cell apoptosis was then analyzed by flow cytometry (BD FACS Calibur, San Jose, CA) with BD Accuri C6 software.

### Cell cycle assay

A549 cells were seeded in 12-well plates. After 24 h of culture, the plasmids or miRNA mimics were transfected into cells for 24 h. The cells were then detached, washed twice with PBS, and fixed in 70% ethanol overnight. The fixed cells were washed and resuspended in 500 μL PBS containing RNase (50 μg/mL) at 37°C for 30 min. Afterwards, the cells were stained with PI (20 μg/mL) for 30 min, and PI intensity were measured by flow cytometry ((BD FACS Calibur, San Jose, CA).

### Animals and *in vivo* bioluminescence imaging

All animal studies were performed according to the Guidance for the Care and Use of Laboratory Animals approved by Xidian University. Six-weeks female Balb/c mice were purchased from the Chongqing Tengxin Biotechnology Co. Ltd. For *in vivo* imaging, the transfected cells were blown off with PBS buffer, and 1×10^7^ cells were injected subcutaneously onto the thighs of Balb/c mice (n=3). Mice were anesthetized with 2% isoflurane in oxygen, luciferase substrate D-luciferin (150 mg/kg, Yeasen Biotechnology Co., Ltd, Shanghai, China) or coelenterazine (4 mg/kg, Yeasen Biotechnology) was injected intraperitoneally onto mice. 5 min later, Xenogen Lumina Π system (Caliper Life Sciences) was used to acquire luminescence signals. The luminescence intensity was expressed as p/s/cm^2^/sr using the Living Imaging Software 4.1.

### *Ex vivo* blood detection of Gluc activity

For hydrodynamic injection, the miCop-Gluc plasmids (10 μg) were dissolved in 10% normal saline (ml) of the mouse body weight (g), and quickly inject it into the body by through the tail vein of the mouse within 5-7s. 24 h later, 2.5 μL of blood was collected from the tail vein of mice at different time point and mixed with 500 μL of 20 mM ethylenediaminetetraacetic acid (EDTA). 50 μL of coelenterazine substrate was added to the wells and Gluc activity was measured using a Glomax-20/20 luminometer (Promega).

### Statistical analysis

All experiments were performed in triplicate independently. All data are presented as mean ± SD. Student’s t tests were used to analyze statistical significance between experimental groups. P value < 0.05 was considered as statistically significant.

## Acknowledgements

This work was supported by National Natural Science Foundation of China (No. 32271512), Natural Science Basic Research Program of Shaanxi (No. 2022JC-56, 2023-JC-ZD-43), Science and Technology Projects in Guangzhou, China (No. 202206010049).

