## Supplementary Figure S1 for "Engineering synthetic CopT/A-based genetic circuits for miRNA imaging and functional gene regulation"

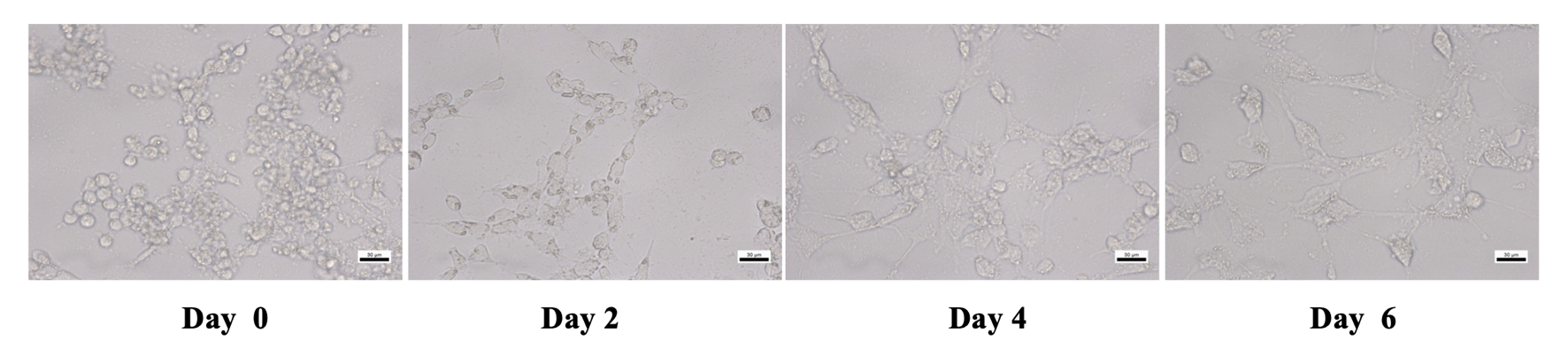

**Supplementary Figure S1. Morphological changes of P19 cells during differentiation.** After P19 cells were treated with RA for different days, the cell morphology was observed by microscope. The protrusions grown by the cells gradually increased and extended, and the P19 cells gradually transformed into neurons.
